# Mapping of multiple complementary sex determination loci in a parasitoid wasp

**DOI:** 10.1101/620229

**Authors:** Cyril Matthey-Doret, Casper J. van der Kooi, Daniel L. Jeffries, Jens Bast, Alice B. Dennis, Christoph Vorburger, Tanja Schwander

## Abstract

Sex determination has evolved in a variety of ways and can depend on environmental and genetic signals. A widespread form of genetic sex determination is haplodiploidy, where unfertilized, haploid eggs develop into males and fertilized diploid eggs into females. One of the molecular mechanisms underlying haplodiploidy in Hymenoptera, a large insect order comprising ants, bees and wasps, is known as complementary sex determination (CSD). In species with CSD, heterozygosity at one or several loci induces female development. Here, we identify the genomic regions putatively underlying multi-locus CSD in the parasitoid wasp *Lysiphlebus fabarum* using restriction-site associated DNA sequencing. By analysing segregation patterns at polymorphic sites among 331 diploid males and females, we identify four CSD candidate regions, all on different chromosomes. None of the candidate regions feature evidence for homology with the *csd* gene from the honeybee, the only species in which CSD has been characterized, suggesting that CSD in *L. fabarum* is regulated via a novel molecular mechanism. Moreover, no homology is shared between the candidate loci, in contrast to the idea that multi-locus CSD should emerge from duplications of an ancestral single-locus system. Taken together, our results suggest that the molecular mechanisms underlying CSD in Hymenoptera are not conserved between species, raising the question as to whether CSD may have evolved multiple times independently in the group.

**Author summary:** The genetic or environmental signals that govern whether an organism develops into a male or female differ across species, and understanding their evolution is a key aspect of biology. In this paper, we focus on complementary sex determination (CSD), a genetic sex determination system found in many species of bees, ants and wasps where heterozygosity at one or multiple genetic regions determines the sex of the individual. We identify multiple genetic regions in the parasitoid wasp species *Lysiphlebus fabarum* that are likely underlying CSD. We show that these candidate CSD regions share no similarity with each other and that they differ from the CSD region known in the honey bee, the only species with a well-characterized CSD system. Our results suggest a different molecular mechanism underlying CSD in the wasp and that multiple CSD regions do not necessarily arise from duplications as generally thought.

## Introduction

A common mechanism of sex determination in animals is via genetic factors, for example by sex chromosomes or sex-specific ploidy [1]. Haplodiploidy is a widespread genetic sex determination system found in approximately 12% of all animal species [1], encompassing some groups of beetles and mites, whiteflies, as well as the whole insect orders Thysanoptera (thrips) and Hymenoptera. In haplodiploid sex determination, unfertilised eggs develop into (haploid) males, and fertilised eggs develop into (diploid) females. In many haplodiploid hymenopteran species, the molecular mechanism underlying female development depends on heterozygosity at the complementary sex determination (CSD) locus [2, 3]. Female development is induced when the individual is heterozygous at the CSD locus, and male development is induced for individuals with only one allele at the CSD locus (either via homo- or hemizygosity). In the honeybee, the only organism where the CSD locus has been characterized so far, *csd* is a paralog of *transformer*, a key gene in the sex determination pathway of insects [4]. The precise mechanism by which CSD regulates sex determination is still unknown, but it is believed that the formation of a heterodimer is key to triggering the female developmental pathway [5].

CSD-based sex determination generates a significant genetic load under inbreeding, as low allelic diversity results in the production of CSD-homozygous, diploid eggs. Depending on the species, the resulting diploid males can have reduced fertility and/or survival [6]. It is thought that multilocus CSD (ml-CSD), a derived mechanism, has been favored under these conditions [7, 8]. In species with ml-CSD, female development is induced if at least one of the CSD loci is heterozygous. Thus, haploid eggs develop into males, as they are hemizygous for all loci, and diploid males are only produced if individuals are homozygous at all loci [9].

CSD loci can be found by identifying genomic regions for which females are heterozygous and diploid males are always homozygous. In many species, diploid males are difficult to come by, because of their reduced fitness [3]. However, we uncovered many diploid males while studying asexual reproduction (thelytokous parthenogenesis) in the parasitoid wasp *Lysiphlebus fabarum* (Hymenoptera: Braconidae), providing a rare opportunity to identify the CSD loci in this species. *L. fabarum* has both sexual and asexual lineages, and CSD is thought to consist of multiple loci, although the actual number of loci remains unknown [10]. In asexual *L. fabarum* the cytological mechanism underlying thelytokous parthenogenesis is central-fusion automixis [11], which involves meiosis followed by a secondary restitution of diploidy through fusion of two meiotic products originating from homologous chromosomes. In this form of automixis, transitions to homozygosity and the associated production of diploid males happens in regions distal to recombination events (Fig 1a). Asexual production of females therefore predicts that at least some CSD loci are close to the centromeres, where heterozygosity is maintained in the long-term (Fig 1b) [12].

**Fig 1.**
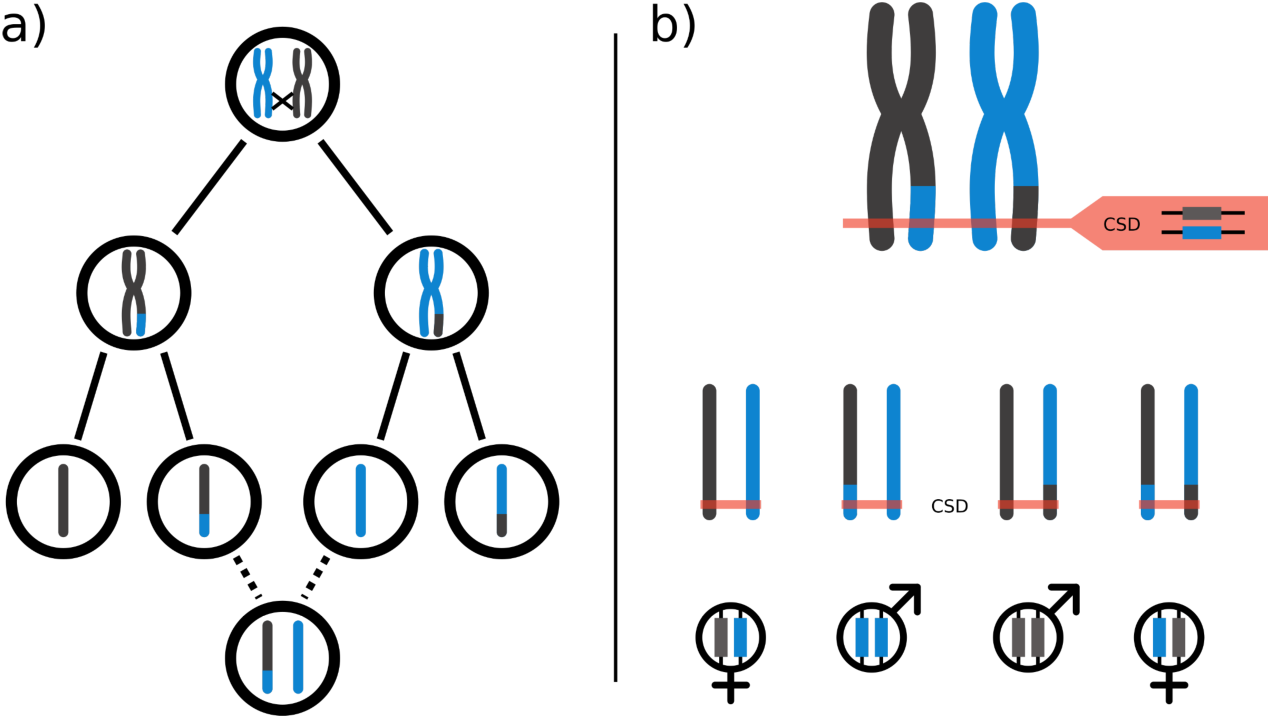
Central fusion automixis and CSD. a) Parthenogenesis with central fusion automixis and crossing over. Homologous chromosomes from the mother are represented in grey and blue respectively. The oocyte undergoes normal meiosis, until two meiotic products originating from homologous chromosomes fuse to form a diploid egg. Chromosomal regions distal to a recombination event become homozygous. b) Interaction between central-fusion and CSD. Visual representation of possible CSD genotypes in the case of a focal CSD locus distal to a recombination event. Assuming a recombination event occurred between the centromere and the CSD locus, an egg produced by central-fusion has a 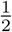 chance to develop into a diploid male. The proportion of recombinant offspring at random loci, should converge towards 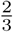 as the number of crossing over increases, thus the chance for a heterozygous locus to become homozygous tends to 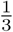 as the number of recombinations increase (see [10], appendix A).

In this study, we explore the genetic basis of CSD in *L. fabarum* using 331 diploid males and females generated in a laboratory cross. Using restriction-associated DNA sequencing (RADseq) [13] and association mapping, we identify regions that are highly homozygous in diploid males, and heterozygous in females. We identify four candidate CSD regions, all on different chromosomes and of which three are close to putative centromeric regions. These loci feature no homology to each other or the known CSD locus in the honeybee, suggesting that the molecular mechanism underlying haplodiploid sex determination is different in *L. fabarum* than in species studied so far.

## Results

### Samples and sequencing

A crossing experiment designed to study the genetic basis of asexuality in *L. fabarum* (Methods, Fig S1) generated 45 families consisting of virgin asexual mothers with (diploid) daughters and both haploid and diploid sons. We genotyped 569 individuals of the 45 families by restriction-associated DNA-sequencing (RADseq), via aligning to an available *L. fabarum* reference genome (see methods for details). After excluding 42 individuals with poor alignment statistics (<10% aligned reads compared to the average of all samples), we used STACKS (version 1.48) to generate a SNP catalog from the 527 remaining samples (380 males, 147 females; see methods for details).

In addition to diploid males and females, asexual females produce haploid males. Such vestigial (haploid) male production is fairly common in asexuals [14, 15]. Since our approach relies on the comparison of heterozygosity between diploid males and females, haploid males are not informative. Because haploid and diploid males are phenotypically identical in *L. fabarum*, we used 899 high confidence SNPs with a minimum sequencing depth of 20 to distinguish them. We considered all males with more than 90% homozygous polymorphic sites as haploids (Fig S2). Using this strategy, we removed 196 haploid males from the dataset and kept 184 diploid males for further analyses.

### Identification of CSD regions

After excluding haploid males, SNP calling was performed again on the 331 diploid individuals (males and females) with a more permissive sequencing depth filter (Methods), yielding 1,195 SNPs (corresponding to an expected density of one SNP every 65kb on the 140Mb *L. fabarum* genome). On average, each sample presented 1,143 SNPs with a mean coverage of 100X. Of the 1,195 SNPs, 723 (61%) were on contigs anchored on chromosomes and were used in subsequent analyses. To identify CSD candidate regions, we then used a case-control design where we test the association of allelic state (homozygous and heterozygous) with sex (male and female). For each SNP, we performed a Fisher-exact test to calculate a score based on the relative proportion of homozygous males versus females. High scoring SNPs are therefore consistently found in a heterozygous state in females and/or in a homozygous state in males. After multiple testing correction (FDR=10^−5^), we found four candidate regions, all on different chromosomes (Fig 2), among which two, on chromosomes 3 and 5, are highly significant (p<10^−6^). Estimated recombination rates between separate SNPs in each region were homogeneous, suggesting the presence of a single CSD locus per region (Fig S3).

**Fig 2.**
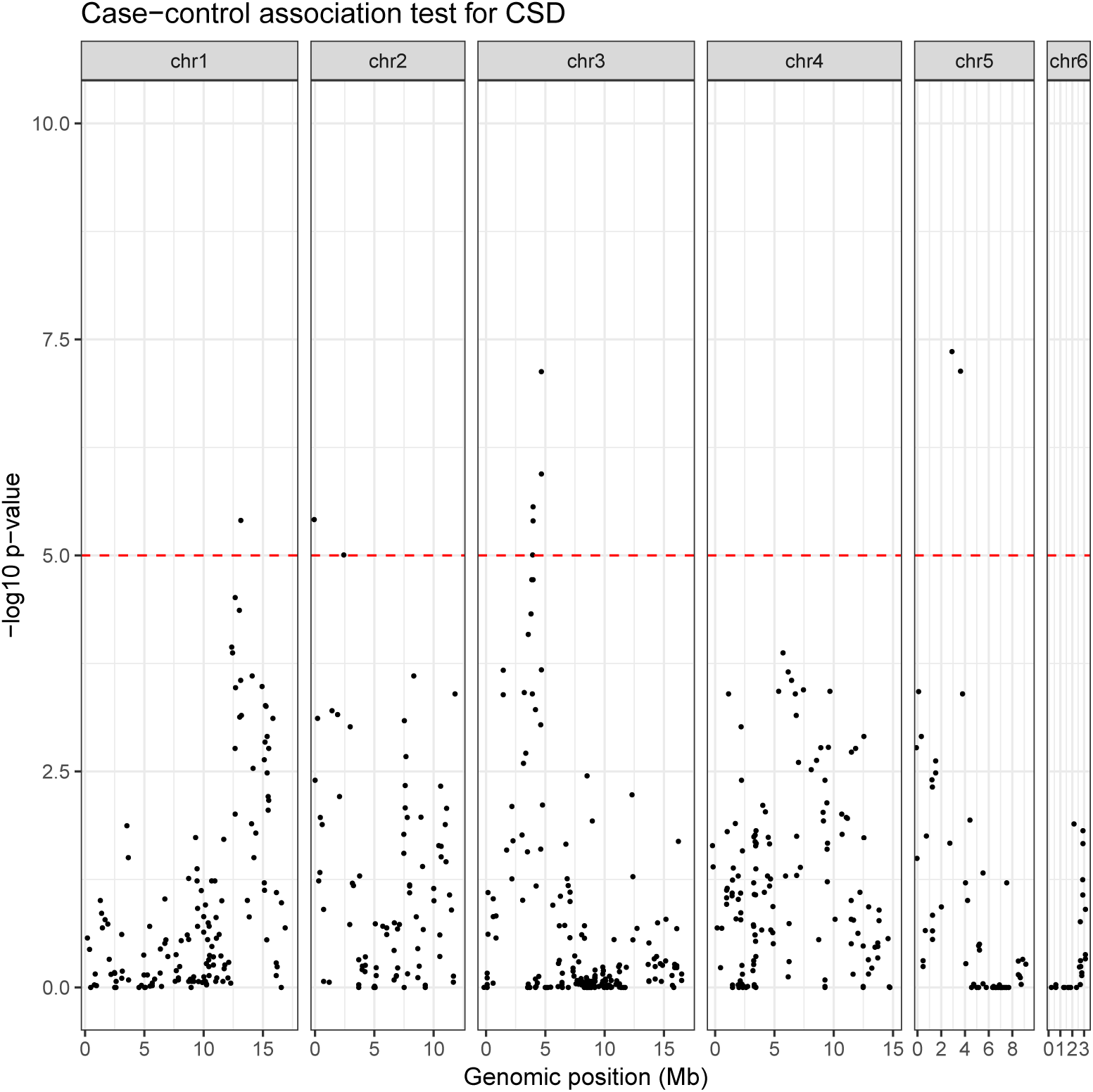
Four CSD candidate regions in *L. fabarum*. Association mapping using one-sided Fisher exact tests for identifying SNPs with an excess of heterozygotes in females relative to males. The Manhattan plot shows -log10 p-values, after Benjamini-Hochberg correction for multiple testing. The different panels show data for different chromosomes, with the horizontal red dashed line showing the p = 10e-5 threshold.

We further assessed the fit of each candidate region to expected heterozygosity levels of CSD regions. According to the CSD model, diploid males must be homozygous at all CSD loci, whereas females need only be heterozygous at one locus. Candidate regions on chromosomes 1, 2 and 3 show a very low proportion of heterozygous males, while this proportion is much higher on the region from chromosome 5 (Fig S3), meaning that the support for the CSD candidates on chromosomes 1, 2 and 3 is stronger than for candidates on chromosome 5.

We also attempted to use polymorphism levels, which are expected to be high for CSD loci, as an additional approach to compare CSD candidate regions. Diploid males having reduced fitness, rare CSD alleles are under positive selection in the wild, leading to balancing selection at CSD loci [16]. Regions undergoing such balancing selection should show elevated levels of nucleotide diversity in wild populations. This was shown to be the case for the CSD locus of the honeybee [17]. We quantified diversity levels across the genome in *L. fabarum* using whole genome sequencing data from 15 individuals collected in a natural population, but we did not detect a significant rise in diversity around any of the candidate regions (Fig S4). This might be explained by much weaker balancing selection on each individual CSD locus under ml-CSD as in *L. fabarum* than on the one locus in sl-CSD species such as the honeybee.

### Location of centromeric regions

As CSD regions are expected to be close to centromeres in asexual *L. fabarum* [10], *we identified the most likely location of the centromere on each chromosome of the L. fabarum* genome. Under central-fusion automixis, the genotype of any diploid offspring should be identical to that of their mother, except for regions distal to recombination events that can become homozygous (Fig 1); this causes homozygosity to increase with distance from the centromeres. Thus, for a locus that is heterozygous in the mother, the proportion of homozygous offspring (male or female) can be used as a proxy for the distance of that locus to the centromere.

Using this approach, we could clearly infer the likely location of the centromeric regions for five out of six chromosomes (Fig S5). Modeling recombination rates using both weighted local regression and moving averages yielded similar results (Fig S5). For the sixth chromosome, which has a smaller number of markers, it was impossible to make a reliable inference of the centromeric region. Out of the four candidate CSD regions, three are close to the estimated centromere locations (chromosomes 1, 3 and 5, Fig 3). Such proximity is expected in organisms with central-fusion automixis, as CSD loci that are further from centromeres would be rendered homozygous in case of recombination [12], causing the loss of their heterozygosity-dependent feminizing effect.

**Fig 3.**
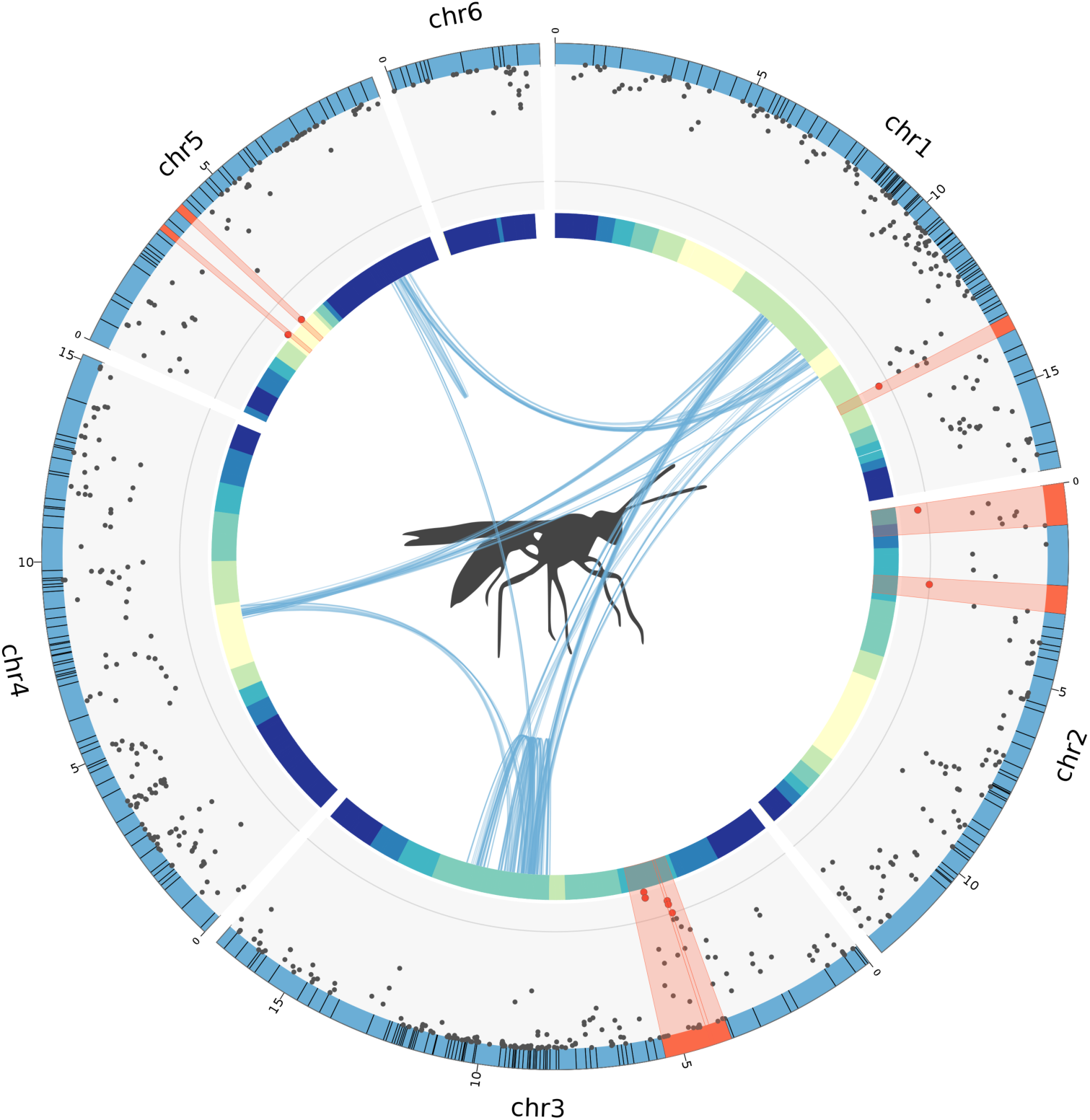
Position of CSD candidate regions relative to centromeres and collinearity blocks. This plot integrates three layers of information from the current study. Each blue segment forming the outer circle represents a chromosome and the black lines intersecting them illustrate the boundaries of anchored contigs. The scatterplot shows the Manhattan plot from figure 2 with highly significant (p <10^−3^, grey line) SNPs and their corresponding contigs shown in red. The inner colored circle is a heatmap showing the loss of heterozygosity (likely location of the centromere) along chromosomes from low (yellow) to high (blue), estimated from proportion of recombinant offspring (Fig S5). Blue curves in the middle show collinearity blocks obtained using MCScanX with default parameters.

### Collinearity across CSD regions

The molecular mechanisms underlying ml-CSD have not been studied thus far, but a verbal model suggested that ml-CSD may derive from a sl-CSD system via duplication of the original CSD locus [8, 18]. We therefore evaluated whether the multiple CSD regions of *L. fabarum* could have evolved via duplication. A common approach to infer gene duplications across different genomic regions is to look for collinearity; the conserved order of homologous genes between regions. We used MCScanX [19] to investigate genome-wide patterns of collinearity in genes and transcripts. The coordinates used were generated by combining gene tracks from the MAKER annotation pipeline [20] and coordinates from aligned transcripts from larvae [21] (SRA accession numbers SAMN10024115-SAMN10024165). We found no evidence for collinearity between candidate CSD regions, suggesting the different CSD loci in *L. fabarum* did not evolve via duplication (Fig 3). A genuine absence of collinearity could mean either that the genetic elements differ across loci, or that the similar region is not large enough to be detected using collinearity. It is also possible that the assembly is too fragmented to detect collinearity, with unanchored contigs interrupting collinearity blocks inside chromosomes. Indeed, the genome is split into 1698 contigs of which 296, accounting for 53.5% of the assembly length, were anchored to chromosomes using a linkage map (Table S2). However, this should not prevent us from identifying a paralogy across CSD regions, as the association mapping revealed very few high scoring SNPs in unanchored contigs (Fig S6).

Improving the placement of contigs or the genome assembly in future studies will allow to reduce the technical constraints for detecting paralogy. Nonetheless, the association mapping step is not affected by genome completeness and identified CSD candidate regions lay the foundations for more detailed molecular characterization of each region.

### Transformer homology

The upstream molecular mechanisms underlying CSD in *L. fabarum* are likely different from those in the honeybee. In the honeybee, which is the only system where CSD has been studied in detail [22], the *csd* gene derives from a duplication of *transformer* (called *feminizer* in the honeybee), a major sex determination gene in insects [23]. The honey bee *csd* gene has been under strong positive selection, resulting in its neo-functionalization as the master switch for sex determination. To investigate whether a homolog of the *transformer* gene is present in the *L. fabarum* candidate CSD regions, we retrieved the protein sequences of *transformer* homologs from eight different hymenopteran species and searched the *L. fabarum* genome using TBLASTN. This approach allowed us to identify a *transformer* homolog on chromosome 1 at 7Mb (positions: 7,006,657-7,006,839), but there was no homolog in any of the candidate CSD regions. There was also no additional *transformer* homolog in the unanchored contigs. The absence of *transformer* homologs in the *L. fabarum* CSD regions suggests that the CSD in *L. fabarum* is based on different molecular mechanisms than in the honeybee. As we were able to identify a *transformer* homolog elsewhere in the genome, our results are unlikely due to the *L. fabarum transformer* sequence being too diverged for identification via homology searches(e.g., [24])

## Discussion

We studied the complementary sex determination (CSD) system in the parasitoid wasp *L. fabarum* and identified four candidate CSD regions on different chromosomes. The absence of a transformer homolog in any of these regions suggests a novel molecular mechanism underlying CSD in *L. fabarum*, with an upstream cue that differs from the one in the honeybee. The other non-honeybee species with genomic candidate regions for CSD, the ant *Vollenhovia emeryi*, possesses two *transformer* copies in one of the candidate regions, which led to the suggestion of a conserved CSD mechanism across ants and bees [25]. Our findings suggest that such conservation does not extend to braconid wasps, a clade that diverged from ants and bees approximately 200 million years ago [26].

Our findings suggest that CSD in *L. fabarum* is based on up to four separate loci, in line with previous inferences based on high variation of diploid male production among different lines of asexual females [10]. However, the exact number of different CSD loci in *L. fabarum* remains speculative. For example, there could be additional CSD loci with polymorphism in wild populations but monomorphism in our studied crosses. Furthermore, the level of support varies among the four loci identified in our study. The candidate locus on chromosome 3 is supported by the highest number of SNPs and shows the most significant association between heterozygosity and sex (p <10^−7^). By contrast, the candidate region on chromosome 5 is highly heterozygous in females, but a high proportion of males is also heterozygous, making it a less promising candidate, as males should be homozygous at all CSD loci. This genetic region could, for example, contain genetic factors unrelated to CSD, but be potentially lethal to females when present in a homozygous state, while having no particular effect on males. Identifying the exact number of separate CSD loci in *L. fabarum* is a challenge for future studies.

Ml-CSD should be favored in species with asexual reproduction via automixis (as in *L. fabarum*) or high inbreeding, where it would decrease the load caused by diploid male production [12]. Our laboratory cross was designed to generate new asexual strains via introgression of asexuality-causing alleles into the genetic background of a sexual species. As a consequence, centromere regions that would never become homozygous under asexuality were rendered homozygous via inbreeding in the sexual generations, resulting in the frequent production of diploid males by the new asexual strains. In wild asexual populations however, loci close to the centromeres will remain heterozygous because of central fusion automixis. CSD loci in asexual populations should therefore be preferentially located in centromeric regions to minimize the production of diploid males. In agreement with this hypothesis, three out of four putative CSD loci we identified are found close to centromeric regions (Fig 3).

Our results call for a reconsideration of the existing theoretical model for the evolution and functioning of ml-CSD. Multi-locus CSD is thought to originate by duplication of an ancestral, single CSD locus [7, 8]. However, the duplication model raises several questions. For example, it does not explain why two recently duplicated CSD loci hemi- or homozygous for different alleles would not be able to complement each other and generate haploid females. Nonetheless such individuals are unheard of in CSD species. In the light of our results, it seems more likely that the different CSD loci have different functions and were not generated via duplication but recruited independently as upstream signals in sex determination. There are currently two known genetic mechanisms underlying haplodiploid sex determination in Hymenoptera [27]: sl-CSD, in the honey bee *Apis mellifera*, and parental genome imprinting, in the jewel wasp *Nasonia vitripennis* [28]. Functional investigation of the CSD regions in *L. fabarum* are likely to reveal a third molecular mechanism of sex determination in Hymenoptera.

## Materials and methods

All scripts and instructions required to reproduce the analysis are implemented in a pipeline available on Github at https://github.com/cmdoret/CSD_Lfabarum. For all analyses, we use contigs from the latest version of the *L. fabarum* reference genome [20] (Lfab v1.0, Available on request: https://bipaa.genouest.org/is/parwaspdb) which have been anchored onto six chromosomes (Table S2) in line with the six chromosomes that were deduced from karyotyping [11]. Raw reads are available on NCBI SRA database under bioproject PRJNA505237, while the anchored genome used here along with all files required to reproduce the analysis are available on Zenodo at DOI 10.5281/zenodo.1488602.

### Samples and RADseq protocol

Samples were obtained from a breeding experiment that was designed to introgress asexuality-causing allele(s) into a sexual line, with the aim to study the genetic basis of asexuality (Fig S1). A haploid male coming from an asexual line of *L. fabarum* was crossed with two females from an iso-female sexual line. Following a crossing design similar to that used in [15], asexual females were obtained in the F3 generation (Fig S1). These asexual females produced diploid sons and daughters, which are the focus of the present study. In total, we used 569 individuals from 45 families, including 11 F3 mothers, 153 F4 daughters and 405 F4 sons. The samples were kept in ethanol at −20°C until DNA extraction using a Quiagen DNeasy Blood & Tissue Kit, following manufacturer instructions. Individuals were sequenced in 6 separate libraries, following the protocol from [29], with the enzymes EcoRI and MseI, and size selection on agarose gel (200-450bp). Samples were multiplexed in each library following the TruSeq multiplexing design, and libraries were pooled by pairs on the same Illumina lane using adapters iA06 or iA12. Single-end sequencing was performed using Illumina Hiseq 2500.

### STACKS pipeline

We used the STACKS software (version 1.48) [30] to process RADseq data. Following quality control using fastqc (https://www.bioinformatics.babraham.ac.uk/projects/fastqc, version 0.11), the raw reads were trimmed and demultiplexed using the “process radtags” module from the STACKS suite and 2 mismatches were allowed to detect adapters. The 93bp trimmed, demultiplexed reads were aligned to the latest assembly of the *L. fabarum* genome using BWA-aln (version 0.7.2) [31] with 4 mismatches allowed. Only uniquely aligned reads were extracted using samtools (version 1.4) [32]. Stacks were then generated from SAM files of unique hits using the Pstacks module, requiring a minimum read depth (-m) of 3 to consider a stack. Individuals with less than 10% uniquely aligned reads compared to the average of all samples were excluded from the analysis. The catalog of loci was built with Cstacks allowing for a distance (-n) of 3 mismatches between samples at each locus. The stacks populations module was run on all samples together, requiring each locus to be present (-r) in at least 80% of samples and have at least a sequencing depth (-m) of 20 for ploidy separation, or 5 for association mapping (Table S1). We also required a minimum allele frequency (–min-maf) of 10%. The different STACKS parameters were selected following guidelines in [33].

### Ploidy separation and filtering

To determine the ploidy of all individuals, we rely on genome-wide homozygosity. Haploid samples are hemizygous and should have extremely high homozygosity levels compared to diploid ones. We included only high-confidence SNPs in the STACKS populations module by using a stringent cutoff of 20 reads for the minimum sequencing depth (-m parameter). 899 high-confidence SNPs passed the quality filters and on average, each sample presented 853 of these polymorphic sites with a (high) mean coverage of 132X. We then computed the proportion of homozygous SNPs per individual using VCFtools (version 0.1.13) [34] on the output VCF file from populations. To account for sequencing errors and paralog merging, a conservative threshold of 90% homozygosity among polymorphic sites was determined empirically based on the bimodal distribution of homozygosity (Fig S2). Individuals above that threshold (n = 196) were considered haploid. All these haploid individuals were males.

Haploid males were used to identify and filter out heterozygous SNPs generated via paralog merging. To this end, we extracted loci that were heterozygous in more than 50% of haploid samples (26 loci) and removed these from the set of loci to analyze in diploid samples. This was done by rerunning populations only on diploids and specifying the list of loci with heterozygous sites in haploids using the blacklist (-B) parameter.

### Association mapping

Case-control association mapping was used to identify CSD-candidate regions, based on the observed heterozygosity at each SNP in males and females. The number of heterozygous males, heterozygous females, homozygous males and homozygous females was computed for every SNP and a one-sided Fisher exact-test was performed for each SNP on the 2X2 contingency table. The alternative hypothesis was that the proportion of homozygous males at the SNP is higher than the proportion of homozygous females. P-values were corrected for multiple-testing using Benjamini-Hochberg correction.

### Centromere identification

The proportion of recombinant offspring per locus along the genome was used to estimate centromere position. In each family, all sites that are heterozygous in the mother were used. If the mother was not available, a site was considered heterozygous if at least one of her offspring was heterozygous or if two offspring were homozygous for different alleles. At each site, the proportion of recombinant (homozygous) offspring was computed among all offspring whose mother was heterozygous (all families pooled). The proportions were then used to visualise recombination rates along the genome using two different methods: 1) computing mean homozygosity in a sliding window containing 30 sites with a step size of 1, and 2) using a weighted local regression model of degree 2 with a span of 0.4 to obtain a smooth estimate curve. The weights given to each site in the local regression correspond to the number of offspring taken into account when computing the proportion of homozygous offspring. For each chromosome, the minimum value of the local regression curve was used to approximate centromere location.

### Nucleotidic diversity

Whole genome sequencing was performed on 15 *L. fabarum* wild samples (11 males and 4 females) using paired-end reads on Illumina HiSeq 3000. The raw reads were trimmed using trimmomatic with LEADING:20 and TRAILING:20 and aligned to the reference genome using BWA-mem with default parameters. SNPs were then called using samtools mpileup (version 1.4) [32] skipping indels, and variants aligning to chromosome-anchored contigs were extracted. *π* nucleotide diversity was computed in sliding windows of 100bp with a step size of 10bp.

### Recombination rates

Recombination rates along the genome were interpolated from the same linkage map that was used to anchor the assembly. Recombination rates are assumed to be uniform between linkage map markers. Thus, the genetic distance between a linkage map marker and a SNP is linearly dependent on their physical distance. Given a SNP S between two linkage map markers M0 and M1 the genetic distance between S and M0 *G*_*M*0*−S*_ is given by 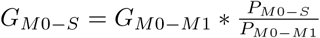 where *P* are physical distances in base pairs.

### Collinearity analyses

Collinearity blocks were defined using the default parameters of MCScanX: A collinearity block is called if two genomic segments share 5 homologous genes in conserved order with at most other 25 genes inserted in between. Gene coordinates were defined by merging maker gene prediction tracks and transcripts assembled from reference-aligned RNAseq reads from 5-days old larvae [21] (SRA accession numbers SAMN10024115-SAMN10024165). Gene sequences were extracted at the merged coordinates using bedtools2 [35] and homologous genes were defined by all versus all blastn, using the blast+ command line tools [36], selecting matches with an e-value inferior to 10e-05.

### Transformer homology

Protein sequences of *feminizer* /*transformer* homologs were retrieved from UniproKB for 8 species of Hymenoptera: *Nasonia vitripennis* (B3VN92), *Euglossa hemichlora* (D9MZ89), *Bombus terrestris* (B4Y115), *Melipona compressipes* (B4XU23), *Apis florea* (A0A0H4URN0), *Apis cerana* (B1NW84), *Apis mellifera* (Q6V5L4), *Apis dorsata* (B1NW85). *Homology with these sequences was searched using TBLASTN against the L. fabarum* genome.

## Supporting information

Supplemental File 1: Genotypes table

Supplemental File 2: Samples description

Supplemental File 3: Gene annotations in CSD regions

## Acknowledgements

We thank Paula Rodriguez for her work with the lab rearings and John Wang for discussions. All computations were performed at the Vital-IT (http://www.vital-it.ch) Center for high-performance computing of the SIB Swiss Institute of Bioinformatics. This study was supported by the Swiss National Science Foundation (http://www.snf.ch), grants PP00P3 139013 and PP00P3 170627 to TS, and SNSF Professorship nr. PP00P3 146341 and Sinergia grant nr. CRSII3 154396 to CV. The funders had no role in study design, data collection and analysis, decision to publish, or preparation of the manuscript. All authors have read and approved the manuscript and it has not been accepted or published anywhere else.

## Supporting information

**Table S1.**
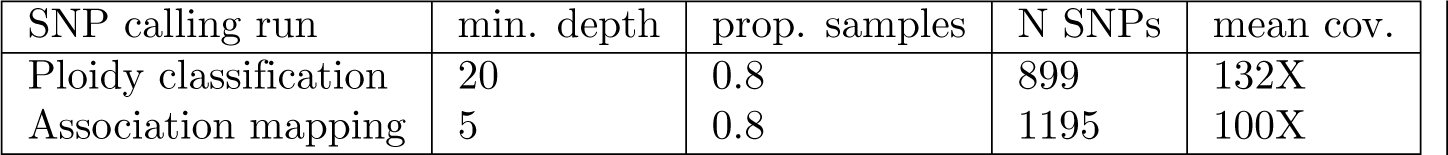
Summary table of SNP calling statistics. The values for STACKS populations filters for minimum sequencing depth (-m) and proportion of samples in which the locus must be present (-r) are shown for both SNP calling runs, along with the output results: Number of SNPs passing the filters, along with mean coverage per individual.

| SNP calling run | min. depth | prop. samples | N SNPs | mean cov. |
| --- | --- | --- | --- | --- |
| Ploidy classification | 20 | 0.8 | 899 | 132X |
| Association mapping | 5 | 0.8 | 1195 | 100X |

**Table S2.** Assembly statistics. Original assembly statistics for the *L. fabarum* genome (Lfab v1.0) and statistics for the subsequent linkage map-assisted anchoring (Dennis et al., in prep.). Genome available on request at https://bipaa.genouest.org/is/parwaspdb.

|  |  |
| --- | --- |
| Assembly length | 140.7Mb |
| Longest contig | 2.2Mb |
| Mean contig length | 82.9kb |
| Median contig length | 30.4kb |
| N50 | 216.1kb |
| Number of contigs | 1698 |
| Anchored bases | 75.3Mb |
| Anchored contigs | 296 |
| Oriented contigs | 73 |
| Unplaced contigs | 1402 |

**Fig S1.**
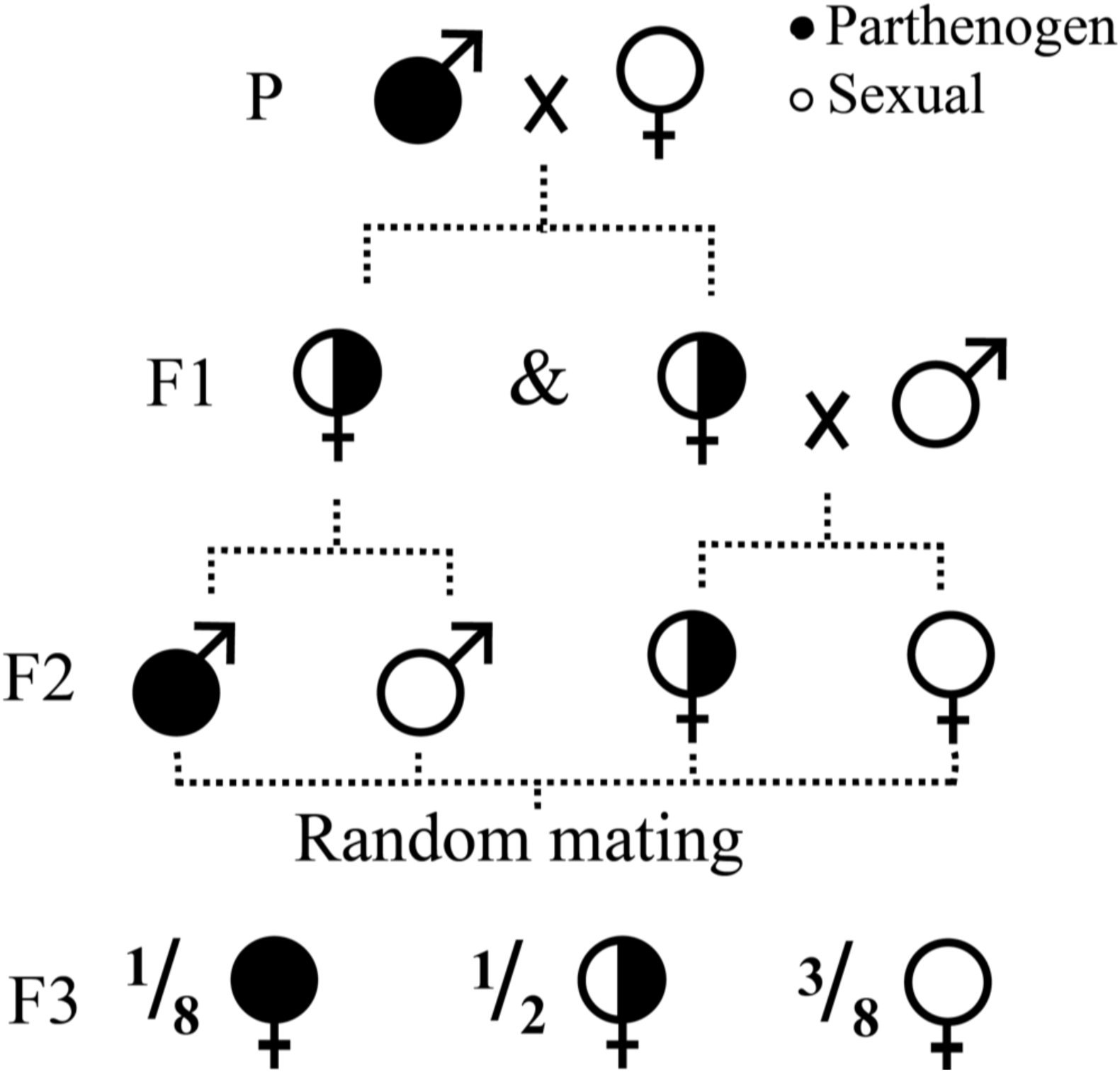
Crossing experiment. Crossing scheme used to establish the families that were included in this study. The asexuality-causing allele is shown in black and sexual phenotypes are shown in white or composite symbols, assuming single-locus recessive inheritance of asexuality (Sandrock et al., 2011) A rare male from an asexual population is mated with a sexual female, yielding F1 females that are heterozygous (the F1 males were discarded). The F1 females were kept isolated to produce recombinant males, or were mated with males from the same sexual population. Following random mating among F2 siblings, the ratio of asexuals to sexuals is approximately 1:7 in the F3 generation. The (diploid male and female) offspring produced by these asexual females were used in the current study.

**Fig S2.**
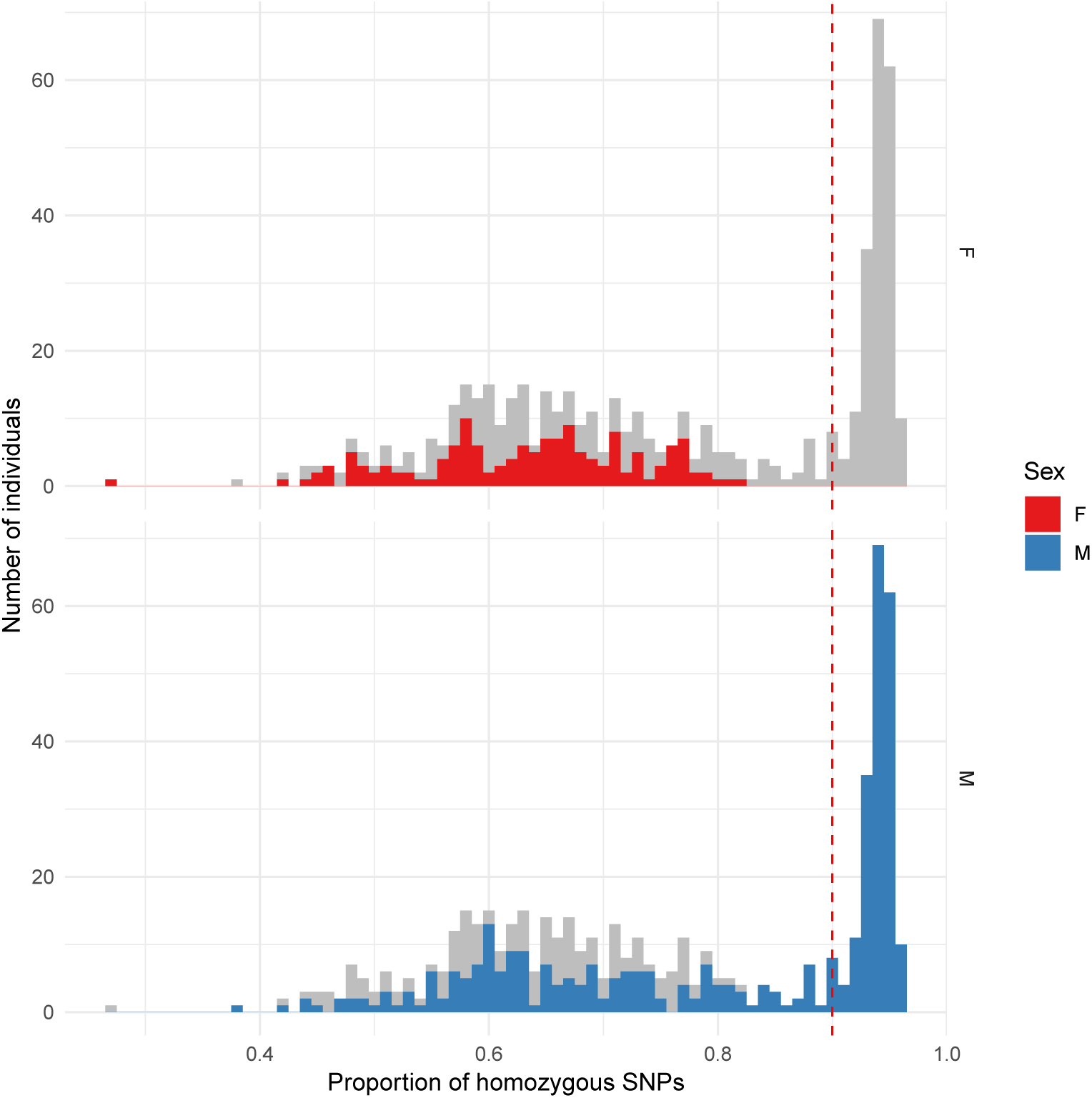
Ploidy classification. Distribution of the genome-wide proportion of homozygous SNPs per individual. Males are represented in blue and females in red. The ploidy classification threshold of 90% homozygosity is shown with a vertical dashed line. Individuals above this value are considered haploid.

**Fig S3.**
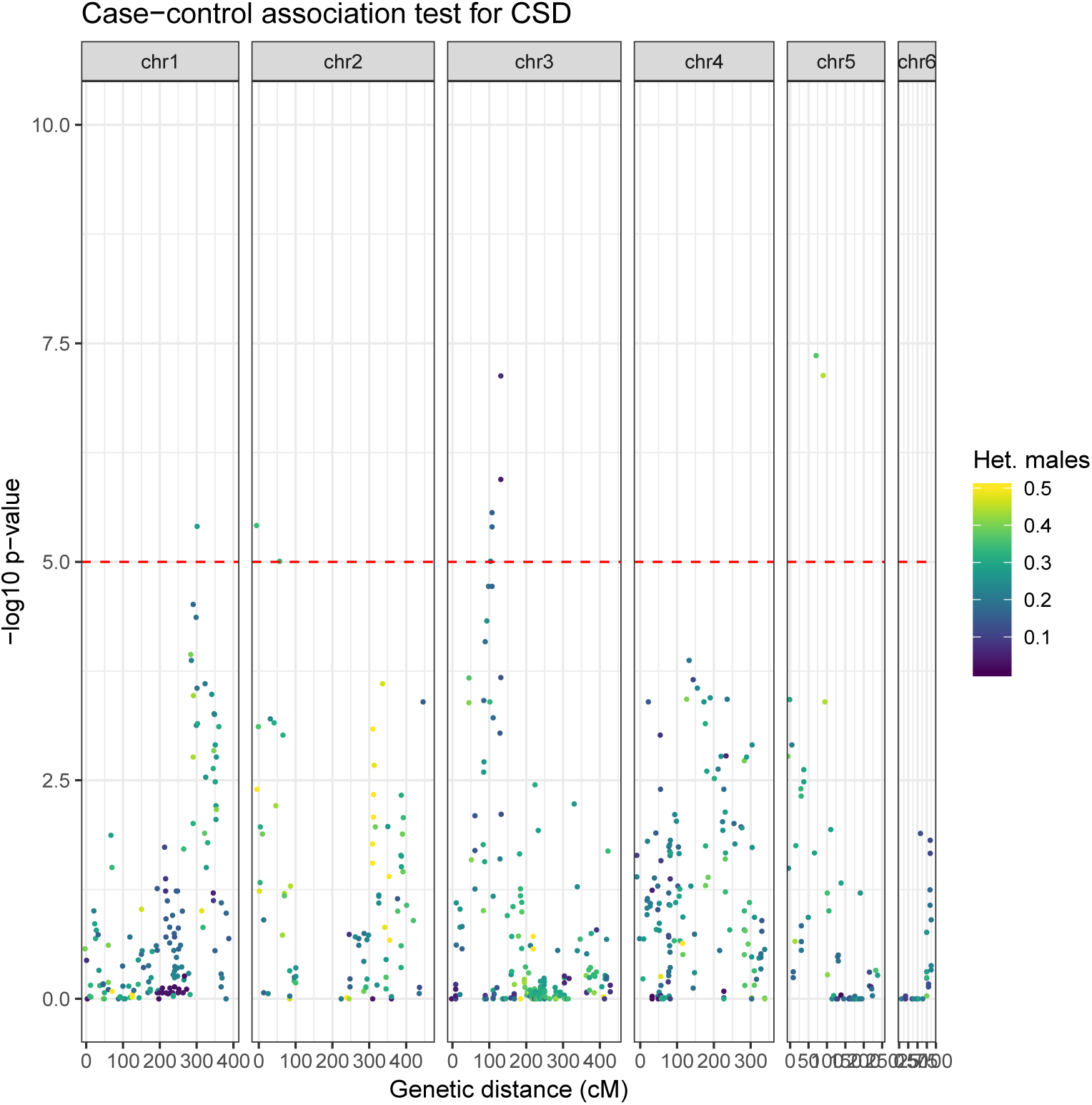
Manhattan plot for the CSD association mapping. SNPs are colored according to the proportion of heterozygous males at the position. Coordinates are in centiMorgans to show genetic distance interpolated from the linkage map.

**Fig S4.**
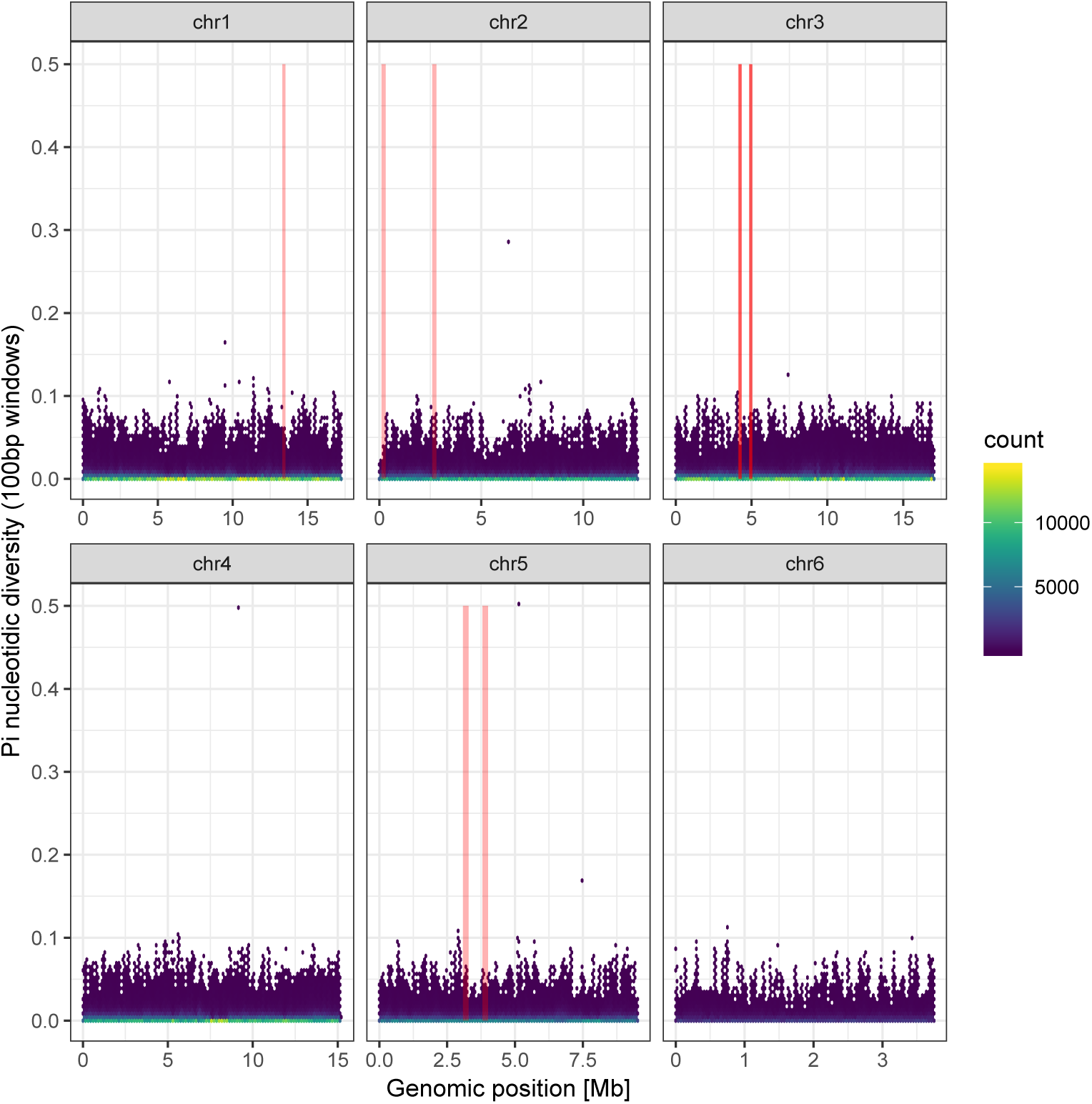
Pi nucleotide diversity along genome. Nucleotide diversity computed in 100bp windows with a step size of 10bp from whole genome sequencing data of 15 wild samples. CSD candidate regions are indicated by red lines.

**Fig S5.**
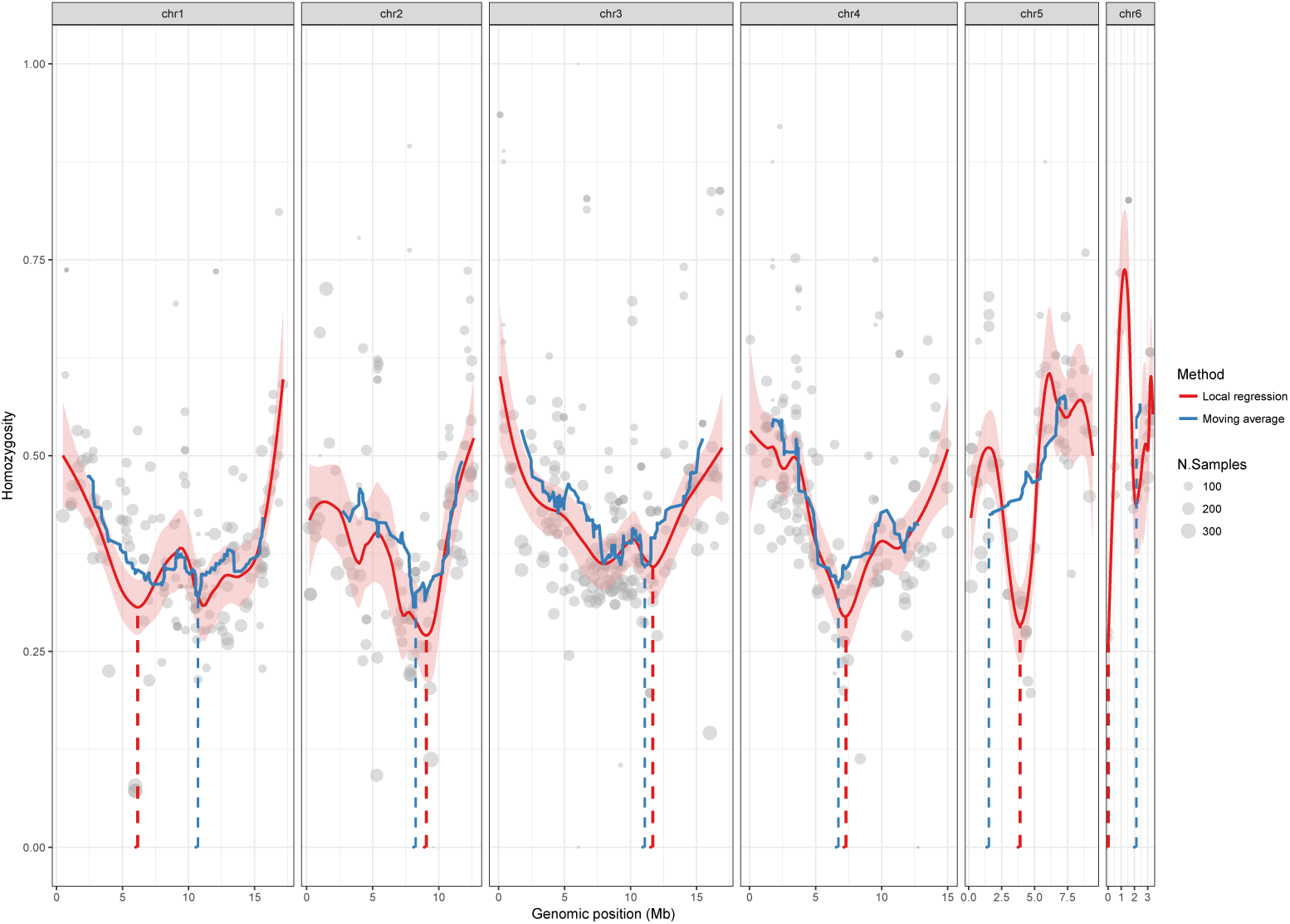
Estimation of centromere locations. Modeling the positions of centromeres from the proportion of recombinant offspring along chromosomes using two independent methods: Weighted local regression of degree 2 and span 0.4 (red) and a moving average with a window size of 30 SNPs and a step size of 1 (blue). 95% confidence intervals of the local regression are shown in light red. The proportion of homozygous (recombinant) offspring at each SNP is displayed as a grey dot, with the area proportional to the number of individuals used to compute the proportion. Each of the six panels represents a chromosome and the vertical dashed lines are the centromeres, defined as the region with lowest proportion of recombinant offspring along the chromosome. Note that since the number of males and females used for centromere identification are similar, the centromere and CSD identifications are not confounded; all samples are expected to be highly heterozygous towards centromeres.

**Fig S6.**
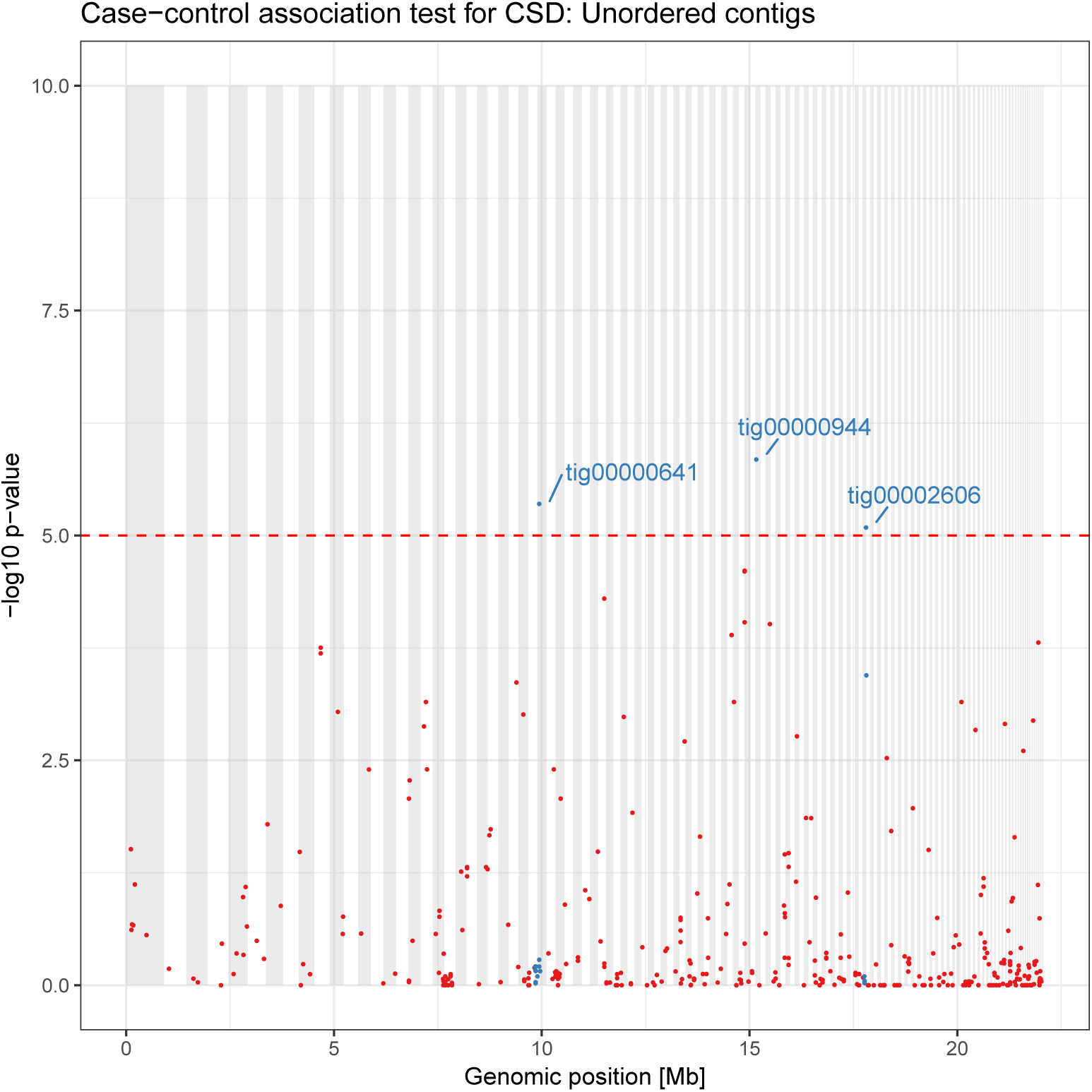
Association scores of SNPs on unanchored contigs, with contigs ordered by size. Log10 Benjamini-Hochberg-corrected p-values from the CSD association mapping for SNPs located in contigs that are not anchored on a chromosome. Alternate grey-white background shows contig boundaries. Significant SNPs are labelled with the name of their contig and all SNPs on the same contig are shown in blue.

**File S1 Genotype table** Genotype table for all individuals at every sites in the RADseq catalog used for the association mapping. Genotypes are encoded as E (Heterozygous), O (Homozygous) or M (Missing).

**File S2 Sample description table** List of all sequenced individuals with their sex, family, generation (F3 being referred to as mothers and F4 as offspring), number of homozygous SNPs observed and expected under Hardy-Weinberg (O(HOM) and E(HOM)), total number of SNPs, the inbreeding coefficient F, mean SNP sequencing depth, proportion of homozygous SNPs, and inferred ploidy (D: diploid, H: haploid). All SNP statistics are computed using vcftools v0.1.15 on the stringent set of loci used for ploidy separation.

**File S3 Gene annotations in candidate regions** List of all predicted gene annotations overlapping with candidate CSD regions. Regions are defined as genomic intervals between significant SNPs (p<10^−5^). Directly adjacent intervals are merged into a single one (i.e. in the case of a single non-sigificant SNP). Blast2go annotations of predicted genes were retrieved from the BIPAA website.

